# GATA2 Expression by Intima-Infiltrating Macrophages Drives Early Atheroma Formation

**DOI:** 10.1101/715565

**Authors:** Charles Yin, Angela M. Vrieze, James Akingbasote, Emily N. Pawlak, Rajesh Abraham Jacob, Jonathan Hu, Neha Sharma, Jimmy D. Dikeakos, Lillian Barra, A. Dave Nagpal, Bryan Heit

## Abstract

Aberrant macrophage polarization is a major contributor to the onset and progression of atherosclerosis. Despite this, macrophage polarization during in early stages of human atherosclerotic disease is poorly understood. Using transcriptomic analysis of macrophages recovered from early-stage human atherosclerotic lesions, we have identified a unique gene expression profile dissimilar to that observed in later stages of disease that is characterized by upregulation of the hematopoietic transcription factor GATA2. GATA2 overexpression *in vitro* recapitulated defects observed in patient macrophages, including deficiencies in the uptake and processing of apoptotic cells, and in the catalysis of atherogenic protein modifications, with GATA2 knockdown abrogating these defects. Our data describe a previously unreported macrophage differentiation state present in early atheroma formation and identifies GATA2 as a driver of macrophage functional defects during the early stages of atherosclerosis in humans.

## Introduction

Macrophages are the primary immune cell type driving the onset and progression of atherosclerosis. Under homeostatic conditions macrophages are atheroprotective, through both the endocytic clearance of lipoprotein deposits in the sub-vascular space, and through efferocytosis – the phagocytic clearance of apoptotic cells. Combined, these macrophage functions prevent the accumulation of lipids and dying cells, and induce anti-inflammatory signaling, thereby countering the pathological processes required for plaque formation^1–4^. Although macrophages are normally capable of processing and exporting cholesterol in large amounts^5,6^, during atherosclerosis the burden of low-density lipoprotein (LDL) and its chemically modified variants such as oxidized LDL (oxLDL) exceed the macrophages’ processing capacity, resulting in the accumulation of intracellular cholesterol and the differentiation of these macrophages into lipid-laden foam cells^7–10^. In response to the cellular stress associated with cholesterol accumulation, foam cells secrete pro-inflammatory cytokines, apoptose, and are cleared by neighbouring macrophages through efferocytosis^11,12^. In parallel, oxLDL signaling through CD36, Toll-like receptor (TLR) 2 and TLR4 on macrophages and other plaque-resident immune cells further exacerbates inflammation^13,14^. Critically, as atherosclerosis progresses, macrophage efferocytosis within the lesion becomes defective and apoptotic foam cells are left uncleared. These apoptotic cells eventually undergo secondary necrosis, generating a necrotic core^15–17^. Additionally, the release of cytokines from plaque-resident immune cells, and alarmins from necrotic cells, induces the recruitment of monocytes which first differentiate into macrophages, and subsequently into foam cells^18,19^. Lysosomal proteins released during necrosis further destabilize the plaque, contributing to atherosclerotic plaque rupture and exposure of thrombogenic factors contained within the plaque, resulting in thrombus formation that may lead to a stroke or myocardial infarct^15,20,21^.

Macrophages are phenotypically plastic cells, whose functional capabilities are determined by their polarization into specialized subtypes. These polarization states are determined by a combination of macrophage ontologeny^22,23^, tissue-specific cues^24^, host age^25^, and environmental cues^26^, which in turn drive transcriptional programs that control macrophage phenotype. These polarization states change in response to alterations in the local tissue microenvironment such as the introduction of inflammatory mediators^26^. Macrophage polarization is tightly entwined with the development of atherosclerosis. In mice, resident aortic macrophages are embryonically-derived, self-renewing cells characterized by the expression of scavenger receptors, MHC II and efferocytic receptors^22^. In mouse models two macrophage sub-types emerge within atherosclerotic lesions: haematopoietically-derived inflammatory (M1) macrophages, and a TREM2^+^ population unique to the atherosclerotic plaque that expresses a mixture of resident-macrophage markers and markers of alternatively activated (M2) cells^27^. While the ontology of human aortic and cardiac macrophages are not well-understood, macrophages with markers of M1-polarization (iNOS, CD86), M2-polarization (MR, dectin-1), and TREM2^+^-polarization (TREM2, CD9^hi^) have been identified in established plaques^27,28^. In advanced stages of disease, other macrophage sub-types may emerge. For example, intraplaque hemorrhage in humans gives rise to the atheroprotective Mhem macrophage sub-type, which scavenge hemoglobin and are resistant to oxidative stress^29,30^. While multiple macrophage polarization states have been identified in mouse models and in human atheroma samples, very little is known of the transcription factors or the tissue- and environment-specific cues which regulate their transcriptional programs. One of the best characterized atheroma-resident macrophage populations is the Mox sub-type, which in mice accounts for 30% of atheroma-resident macrophages^31^. Mox cells are characterized by a transcriptional profile mediated by the transcription factor Nrf2 which, when knocked out, delays atheroma formation^32^. Nrf2 upregulates several genes not normally expressed in M1 or M2 polarized cells, including genes for processing heme, angiogenic factors, and anti-oxidant mechanisms^31^.

Most investigations of macrophage gene expression in atherosclerosis have concentrated on the stages of disease following the formation of fully developed atheromas bearing a necrotic core and fibrous cap (Stage 4/5 or later in the Stary classification system^33^). As such, little is known of macrophage polarization and function during the earlier stages of plaque development. Lipid accumulation and endothelial activation at the lesion drives monocyte recruitment into the plaque and drives M1-polarization via increased recruitment of inflammatory monocytes versus non-inflammatory monocytes^34^. Local proliferation of these monocyte-derived macrophages then further populates the plaque with inflammatory cells^35^. *In vitro* analyses of changes in macrophage gene expression following exposure to atherogenic lipids provides some insight into the transcriptional changes that occur early in disease. Koller *et al*. demonstrated that exposure of RAW264.7 murine macrophages to oxidized lipids resulted in the rapid upregulation of the genes involved in the uptake of oxLDL, apoptosis and cell stress, while genes controlling cholesterol efflux and cell proliferation were downregulated^36^. Using human monocyte-derived macrophages, Ho & Fraiser observed a similar trend following exposure to modified LDL (moLDL), and determined that complement protein C1q accelerates this response^37^. However, these studies were limited to hematogenous macrophages cultured *in vitro*, and therefore lacked the full range of ontological, tissue-specific and environmental signals which drive macrophage polarization and function *in vivo*.

In this study we investigated the transcriptional profile of macrophages recovered from early human aortic plaque. We identified a unique macrophage polarization state notably different from those reported from later-stage disease in both humans and mouse models. This novel differentiation state is characterized by the downregulation of several critical atheroprotective pathways including those involved in efferocytosis and cholesterol processing, and the concurrent upregulation of inflammatory/atherogenic pathways including protein citrullination. Critically, many of these pathways were found to be regulated by the aberrant expression of the transcription factor GATA2, with GATA2 overexpression *in vitro* recapitulating functional defects induced by oxLDL, and shRNA-mediated GATA2 knockdown abrogating these defects. Combined, our data implicate GATA2 as a key regulator of early plaque development in human atherosclerosis.

## Results

### Macrophage Gene Expression in Early-Stage Atherosclerotic Plaque

Human aortic punch samples were obtained from regions of the ascending aorta with minimal macroscopic atherosclerotic disease in patients undergoing coronary artery bypass graft operations. Approximately 75% of punch samples were found to contain evidence of early plaque when stained for histological markers of atherosclerosis (Figure 1A-C) and for macrophages using anti-CD163 (Figure 1D). CD163 was used in lieu of the more common marker CD68, as up to 40% of CD68^+^ cells in plaque are not of macrophage origin^38^. We observed significant levels of CD163 expression across a range of macrophage polarization states *in vitro* and it colocalized well with CD68 in patient plaques (Figure S1). Plaques observed in punch biopsies were characterized by the presence of lipid-laden macrophages that correlated significantly with macrophage infiltration (Figure 1D,E) and diffuse intimal thickening (Figures 1C,F,S2A-C), but lacked evidence of more advanced lesions such as the deposition of extracellular lipids in the intima (Figure 1B,E) or the accumulation of necrotic cells (Figure 1G, S2D). This pattern of macrophage influx and limited lipid accumulation is consistent with early (stage 1 or 2) atherosclerotic plaque development, as per the classification system of Stary *et al*^33^. Macrophages were selectively isolated from CD163-stained tissue sections by laser capture microdissection, producing a cell population with 33-fold enrichment of the monocyte/macrophage-specific marker CD14 relative to the entire tissue section (Figure 1H). For clarity, these early-atheroma macrophages will henceforth be referred to as “intima-infiltrating macrophages”.

**Figure 1:**
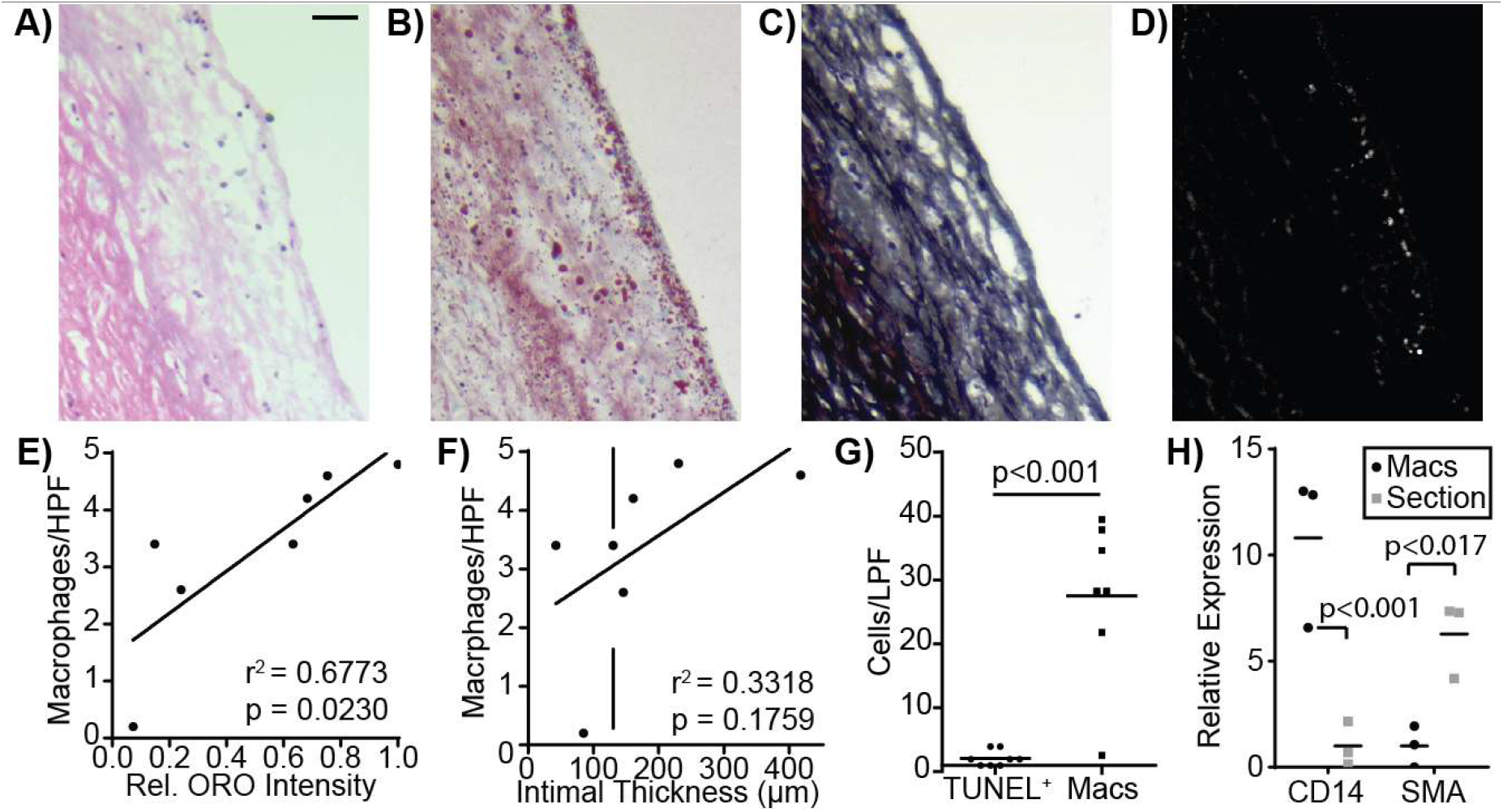
Identification and Recovery of Macrophages from Early Stage Plaque. Regions of plaque development were identified using serial sections from human aortic punch biopsies stained with **A)** H&E, **B)** Oil-Red-O (ORO), **C)** Movats, and **D)** the macrophage marker CD163. **E)** Lipid and sterol deposition, measured by relative ORO staining, correlated strongly with macrophage numbers in high-power fields (HPF, 675 μm × 910 μm). **F)** No correlation was observed between macrophage infiltrates and intimal thickness; vertical line indicates normal aortic intimal thickness. **G)** Number of TUNEL-stained dead cells and macrophages (Macs) in low-power field images (LPF, 2510 μm × 1880 μm). **H)** Fold-enrichment in the macrophage marker CD14 and smooth-muscle marker SMA in purified CD163+ macrophages versus whole tissue sections. Fold-enrichment was calculated using GAPDH-normalized expression levels. Data is representative of (A-D) or quantifies (E-H) a minimum of 5 patients or age- and sex-matched controls. Data is plotted as individual measurements plus either a linear regression (E-F) or median (G-H). p-values were calculated using either linear regression (E-F) or with a Mann-Whitney U test (G-H). Scale bars are 100 μm.

The gene expression profile of these intima-infiltrating macrophages was then compared to M0-differentiated, monocyte-derived macrophages from age- and sex-matched controls, identifying 3,374 differentially expressed protein-coding genes (Figure 2A-B, S3). Principal component analysis demonstrated close clustering of the gene expression profile of intima-infiltrating macrophages between patients, indicating that macrophages in early-stage atheromas develop a consistent gene expression profile (Figure 2C). Gene ontology-based clustering and gene set enrichment analysis were used to identify biological processes that are perturbed in intima-infiltrating macrophages, identifying putative defects in cholesterol homeostasis, the pathways used to engulf and degrade pathogens (phagocytosis) and apoptotic cells (efferocytosis), and antigen processing (Figure S4). As expected, given the large number of differentially regulated genes, multiple transcription factors were significantly up- or down-regulated (Table S1). Of these, we identified GATA2 as the only up-regulated transcription factor that has previously been implicated in coronary artery disease in humans by genetic linkage studies and is known to play an important role in myeloid cell biology.^39–41^ RT-PCR analysis of intima-infiltrating macrophages from five additional patients confirmed upregulation of GATA2 mRNA in intima-infiltrating macrophages (Figure 2D), and immunofluorescence staining determined that GATA2 is predominantly expressed by CD68^+^ cells in this tissue (Figure 2E). GATA2 expression was increased *in vitro* when macrophages were cultured with oxLDL (Figure S5A), and this GATA2 expression could be abrogated by inhibition of the Src-family kinase (SFK)/Syk pathway, but not the ERK1/2 pathway, downstream of the oxLDL receptor CD36 (Figure 2F)^42^.

**Figure 2:**
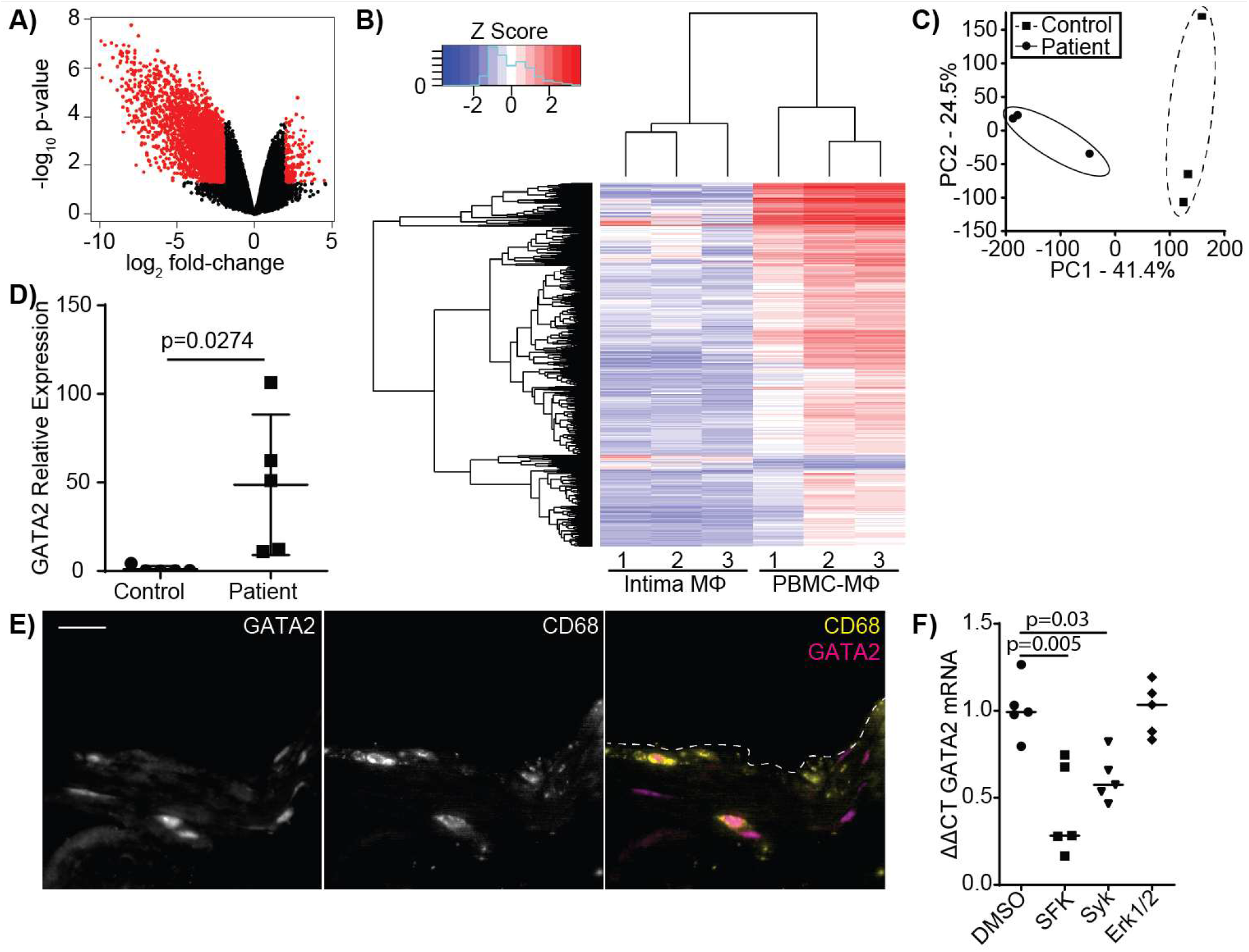
Microarray Analysis of Intima-Infiltrating Macrophages. **A)** Volcano plot illustrating the 3,374 genes up- or down-regulated more than 2-fold in intima-infiltrating macrophages compared to age- and sex-matched monocyte-derived macrophages. **B)** Microarray heatmap showing the gene expression profile of intima-infiltrating macrophages (Intima MΦ) or age- and sex-matched monocyte-derived macrophages (PBMC-MΦ). **C)** Principal component analysis of the gene expression profiles of intima-infiltrating macrophages (Patients) versus Control macrophages. **D)** RT-PCR quantification of GATA2 expression in the intima-infiltrating macrophages versus control macrophages from 5 additional subjects. GATA2 expression is normalized to GAPDH and then normalized to the mean of the controls. **E)** Immunofluorescence image of a representative aortic punch biopsy showing the distribution of GATA2 (magenta) and CD68^+^ macrophages (yellow). Scale bar is 20 μm. **F)** Normalized fold-change in GATA2 expression in THP-1 derived macrophages treated for 72 hrs with 100 μg/ml oxLDL and inhibitors of Src kinases (SFK), Syk and Erk1/2. Data quantifies (A-D,F) or is representative of (E) either 3 (A-C,E) or 5 (D,F) independent experiments. p-values were calculated using a Mann-Whitney test (D) or ANOVA with Tukey correction (F).

### GATA2 Partially Regulates Macrophage Cholesterol Homeostasis

The upregulation of GATA2 in intima-infiltrating macrophages and the dysregulation of multiple cholesterol-processing genes in the same cells suggest that GATA2 overexpression may drive defects in macrophage cholesterol homeostasis. RT-PCR analysis of intima-infiltrating macrophages confirmed that NPC1 and NPC2, which mediate the transfer of endocytosed cholesterol to cytosolic carrier proteins^43^, and ABCA1, which mediates the export of cholesterol from cells^6^, were downregulated in intima-infiltrating macrophages (Figure 3A-C). This alteration in cholesterol uptake and efflux has previously been associated with abnormal cholesterol processing and foam cell development^44,45^. To assess the functional effect of GATA2 expression on cholesterol homeostasis, THP1 macrophages were cultured with oxLDL, which induced a modest time-dependent increase in GATA2 expression (Figure S5A). Because this increase in expression was only 6% of that observed in intima-infiltrating macrophages, we used lentiviral vectors to create THP1 cells expressing GATA2 at levels similar to those observed in intima-infiltrating macrophages, as well as THP1 cells expressing a GATA2 shRNA that blocked oxLDL-induced GATA2 expression (Figure S5B-E). Using this *in vitro* model system, we determined that neither GATA2 overexpression nor the GATA2 shRNA had any effect on cholesterol accumulation (Figure 3D). GATA2 expression was required enhanced cholesterol efflux in normocholesterolemic macrophages, but this effect was lost during hypercholesterolemia (Figure 3E). Combined, these results indicate that cholesterol homeostasis is perturbed in intima-infiltrating macrophages, but this loss of homeostasis is not regulated by GATA2.

**Figure 3:**
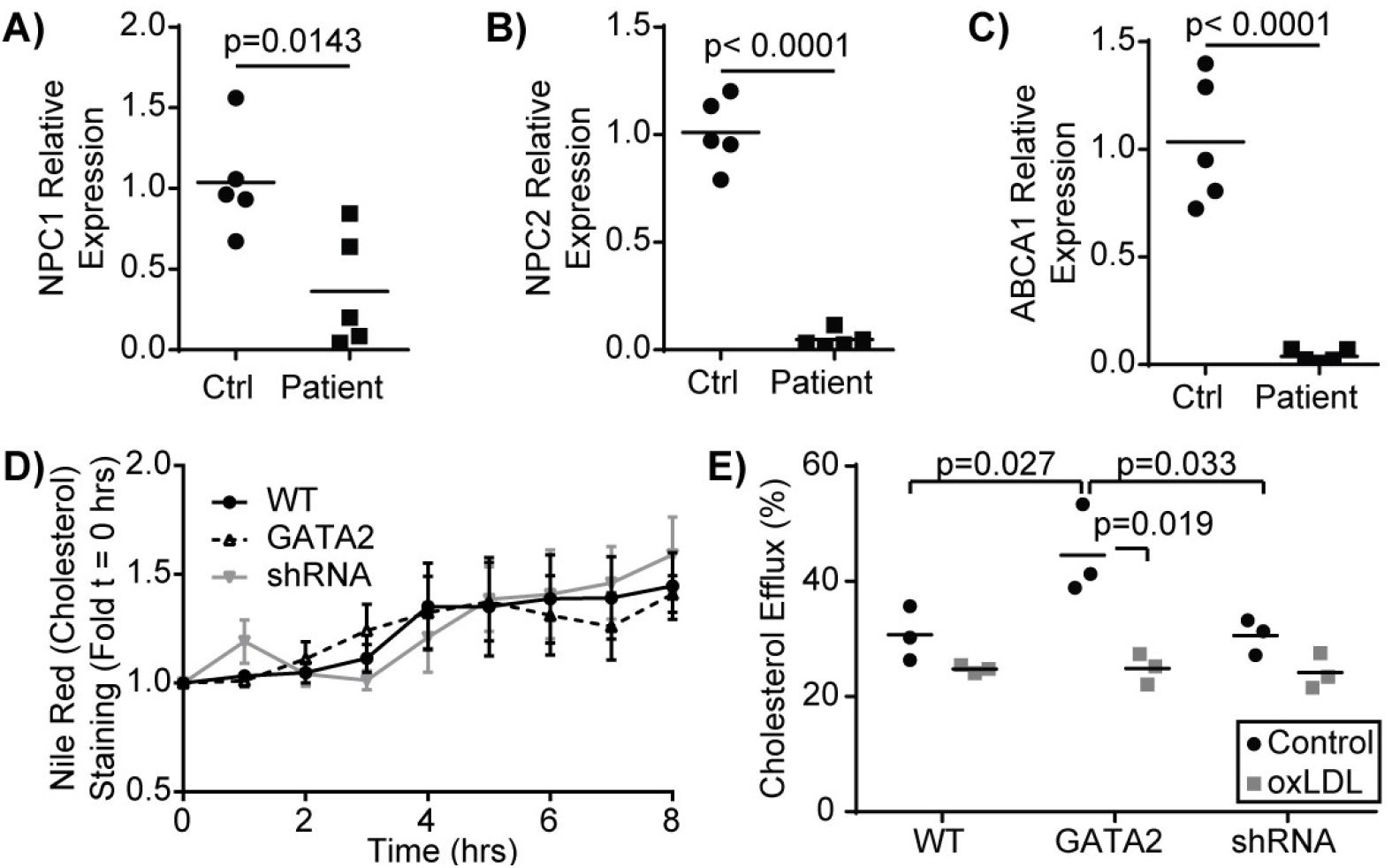
Cholesterol Homeostasis is Partially Regulated by GATA2. RT-PCR was used to quantify the expression of NPC1 **(A)**, NPC2 **(B)**, and ABCA1 **(C)** in intima-infiltrating macrophages (Patients) versus age- and sex-matched monocyte-derived macrophages (Control). **D-E)** Quantification of the effect of expressing either a vector control (WT), GATA2 transgene (GATA2) or GATA2-targeting shRNA (shRNA) on **D)** the accumulation of cholesterol from medium containing 100 μg/ml oxLDL, as quantified by Nile-red live cell microscopy as quantified by foldchange in Nile-red intensity normalized to the time of oxLDL addition (t = 0 min), or **E)** on the effect of normocholesterolemia (Control) or hypercholesterolemia (oxLDL) on the efflux of fluorescently labeled cholesterol. n = a minimum of 3 independent experiments, p-values were calculated using a Students t-test (A-C) or ANOVA with Tukey correction (D-E).

### GATA2 Overexpression Impairs Phagocytosis and Efferocytosis

The efferocytic removal of apoptotic and necrotic cells is a critical atheroprotective mechanism, and failures in this process are a prerequisite for the accumulation of uncleared apoptotic cells which form the necrotic core of advanced atherosclerotic plaques^1,2,46,47^. Our gene expression analysis identified a reduction in the expression of genes involved in the efferocytosis of apoptotic cells and the phagocytosis of microbial pathogens (Figures S4). RT-PCR confirmed that the α_x_ integrin, a receptor for both complement-opsonized pathogens and CD93-opsonized ACs^48^, was downregulated in intima-infiltrating macrophages (Figure 4A). Unexpectedly, integrin-linked kinase, which was downregulated in the patients analyzed in our microarray analysis (Figure S4, Table S1) was not downregulated in the patient cohort used for our RT-PCR assays (Figure 4B). *In vitro*, the phagocytosis of antibody-opsonized particles was profoundly impaired by both culture with oxLDL and GATA2 overexpression. Importantly, this oxLDL-induced defect was abrogated by expression of the GATA2 shRNA (Figure 4C). The same trend was observed in efferocytosis, with both the total number of apoptotic mimics engulfed per macrophage (Figure 4D) and the efficacy of apoptotic cell uptake (Figure 4E) impaired by culture with oxLDL or GATA2 overexpression, and with expression of a GATA2 shRNA restoring normal efferocytic capacity. Combined, these data indicate that GATA2 overexpression impairs the ability of macrophages to recognize and engulf both phagocytic and efferocytic targets.

**Figure 4:**
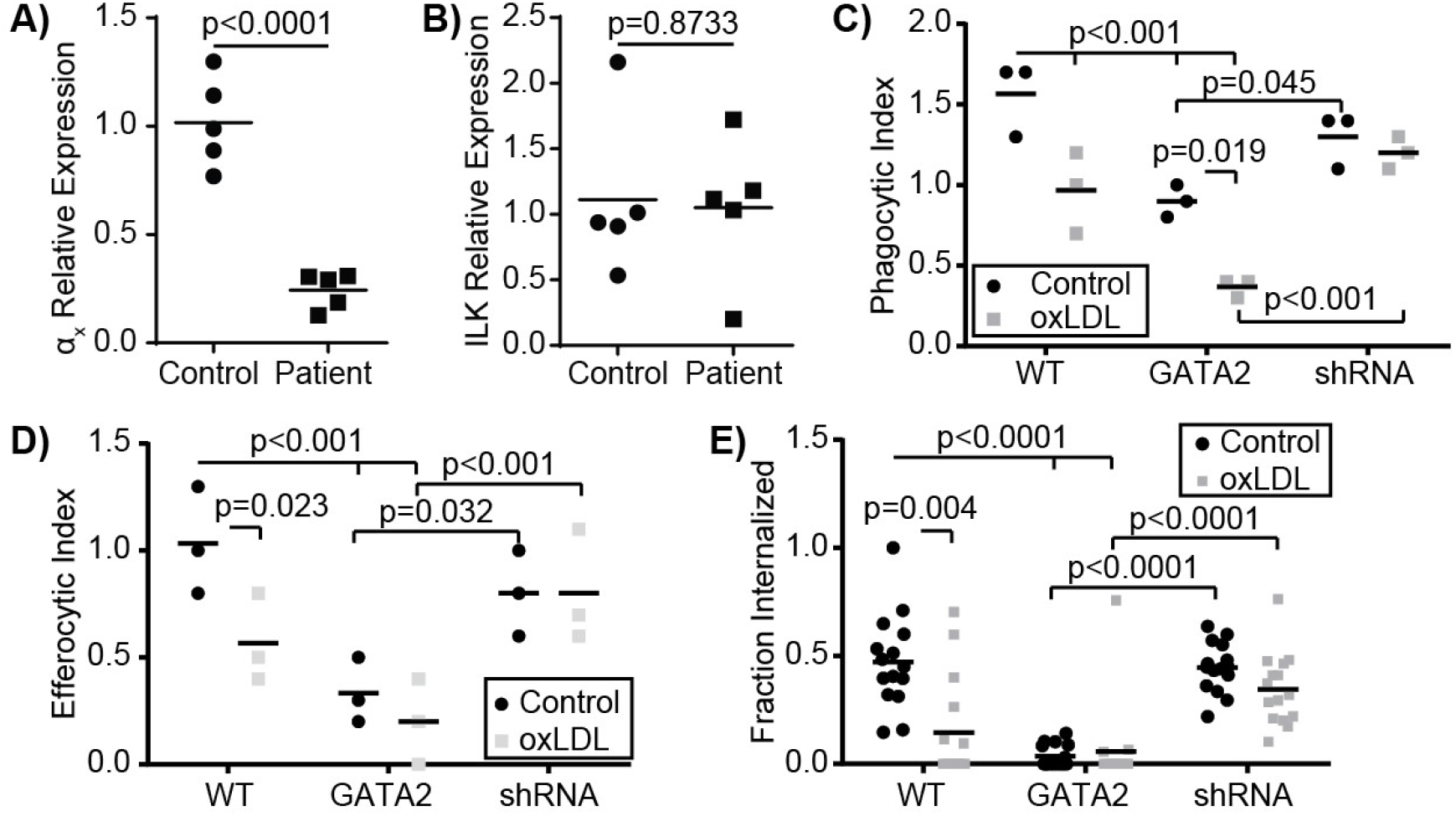
Phagocytosis and Efferocytosis are Impaired by GATA2 Overexpression. RT-PCR was used to quantify the expression of α_x_ integrin **(A)** and integrin-linked kinase (ILK, **B)** in intima-infiltrating macrophages (Patients) versus age- and sex-matched monocyte-derived macrophages (Control). **C-E)** Quantification of the effect of normocholesterolemia (Control) or hypercholesterolemia (oxLDL) on macrophages expressing either a vector control (WT), GATA2 transgene (GATA2) or GATA2-targeting shRNA (shRNA) on **C)** the phagocytic uptake of IgG-coated phagocytic mimics, **D)** the uptake of PtdSer-bearing apoptotic cell mimics, or **E)** the fraction of apoptotic Jurkat cells efferocytosed over 90 min. n = a minimum of 3 independent experiments, p-values were calculated using a Students t-test (A-B) or ANOVA with Tukey correction (C-E).

### GATA2 Overexpression Impairs Phagosome/Efferosome Maturation

In addition to downregulation of receptors and signaling molecules associated with the phagocytic/efferocytic uptake of particulates, intima-infiltrating macrophages also downregulated multiple genes required for the processing of microbes and apoptotic cells following uptake (Figure S4, Table S1). This included down-regulation of Rab7, which is required for phagosome/efferosome fusion with lysosomes^49^, multiple subunits of the vacuolar ATPase which acidifies maturing phagosomes/efferosomes^50^, and two subunits of the NADPH oxidase complex which produces reactive oxygen species to assist in pathogen/apoptotic cell degradation^51^. RT-PCR analysis confirmed the downregulation of Rab7 in intima-infiltrating macrophages (Figure 5A), and phagosome-lysosome fusion defects were observed in oxLDL-treated and GATA2-overexpressing cells, with this impairment prevented by expression of a GATA2 shRNA (Figure 5B,C). In addition to lysosome fusion defects, oxLDL also induced a GATA2-dependent suppression of terminal phagosome pH (Figure 5D). This was not merely a result of poor lysosome-phagosome fusion, as the acidification rate of phagosomes in the first five minutes (i.e. prior to lysosome fusion^52^) was also impaired by oxLDL in a GATA2-dependent manner (Figure 5E). Using the oxidant-sensitive dye nitro blue tetrazolium (NBT), we next determined that, as with lysosome fusion and phagosome acidification, the production of superoxide was impaired by oxLDL in a GATA2-dependent manner (Figure 5F-G). Combined, these data demonstrate that GATA2 overexpression in macrophages causes impairments in multiple processes required for the degradation of targets taken up via phagocytosis and efferocytosis.

**Figure 5:**
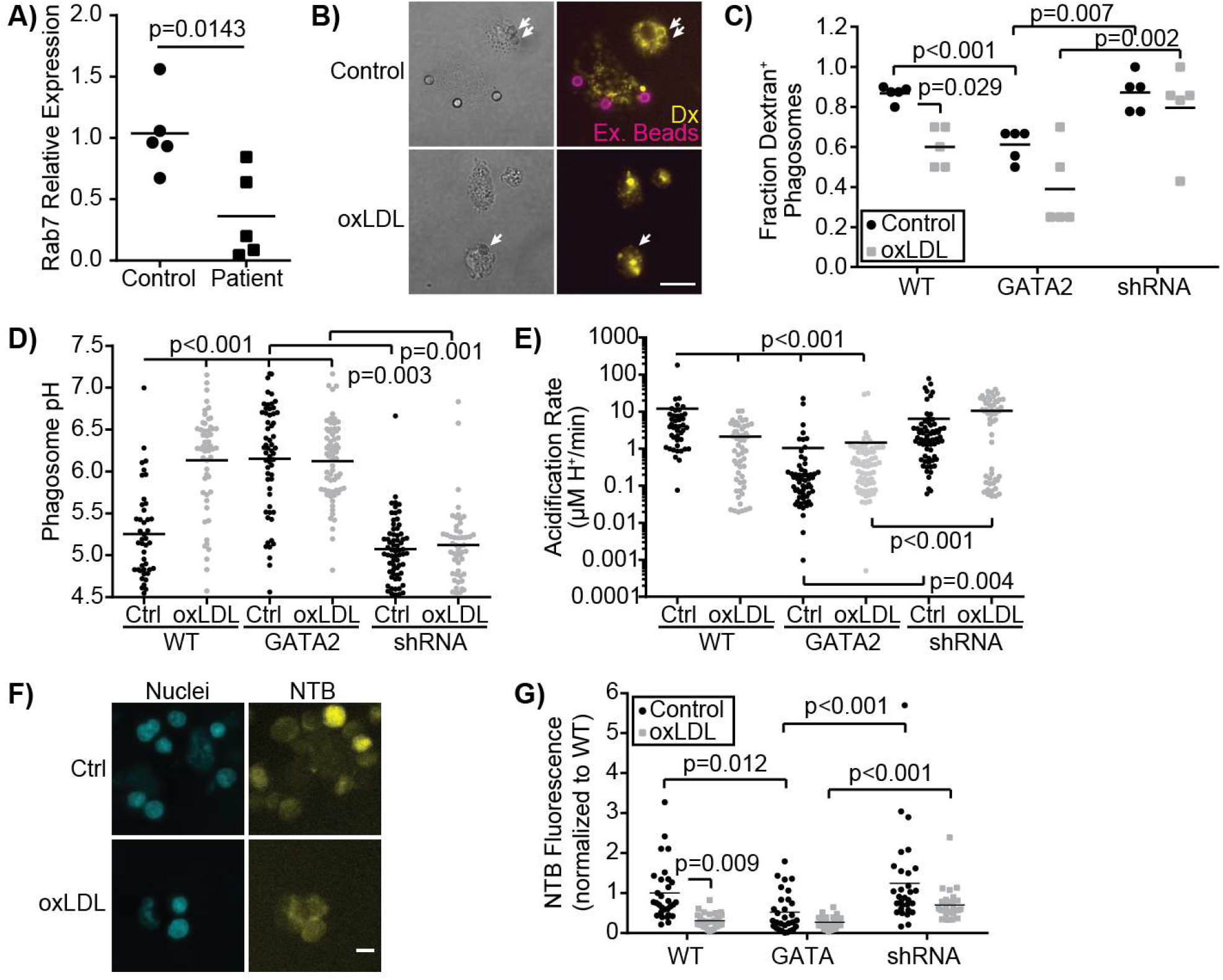
Phagosome/Efferosome Maturation is Impaired by GATA2 Overexpression. **A)** RT-PCR quantification of Rab7 expression in intima-derived macrophages (patient) versus age- and sex-matched monocyte-derived macrophage controls (Control). **B-G)** Quantification of the effect normocholesterolemia (Control/Ctrl) or hypercholesterolemia (oxLDL) on macrophages expressing either a vector control (WT), GATA2 transgene (GATA2) or GATA2-targeting shRNA (shRNA) on **B-C)** the fusion of dextran-loaded lysosomes (Yellow, Dx) with IgG-coated phagocytic targets (arrows), Non-internalized targets are stained with streptavidin-647 (Ex. Beads). Fusion is quantified 30 min after phagocytosis. **D)** Terminal phagosome pH and **(E)** phagosome acidification rate during the first 5 minutes following phagosome closure, as measured using pHrhodo ratiometric imaging. **F-G)** Production of superoxide 90 min after efferocytosing apoptotic Jurkat cells, as measured by NBT fluorescence. p values were calculated using a Students t-test (A) or ANOVA with Tukey correction (C-G, E). Data is representative of (B,F) or quantifies as the ensemble of individual measurements from a minimum of 3 independent experiments. Scale bars are 10 μm (B) or 20 μm (F).

### GATA2 Overexpression Drives Citrullination

We also observed evidence of atherogenic antigen processing in intima-infiltrating macrophages. This included perturbation of the MHC I and MHC II presentation pathways (Figure S4, Table S1), as well as upregulation of enzymes that generate atherogenic self-antigens. This included PADI3, an enzyme which deiminates arginine to form the amino acid citrulline, forming citrullinated self-antigens that contribute to intra-plaque immune complex deposition in some patients^53–55^. RT-PCR analysis confirmed the upregulation of PADI3 in intima-infiltrating macrophages (Figure 6A), and immunohistochemistry identified citrullinated peptides in the intima of these patients’ punch samples (Figure 6B, S6A). Similar patterns of citrullination and PADI3 expression were observed in oxLDL-treated THP1 macrophages (Figures 6C,D, S6B). Using quantitative microscopy, we determined that GATA2 expression is required, but is insufficient, for oxLDL induced protein citrullination (Figure 6E). Together, these data indicate that intima-infiltrating macrophages are engaged in protein citrullination, with this activity regulated in part by GATA2.

**Figure 6:**
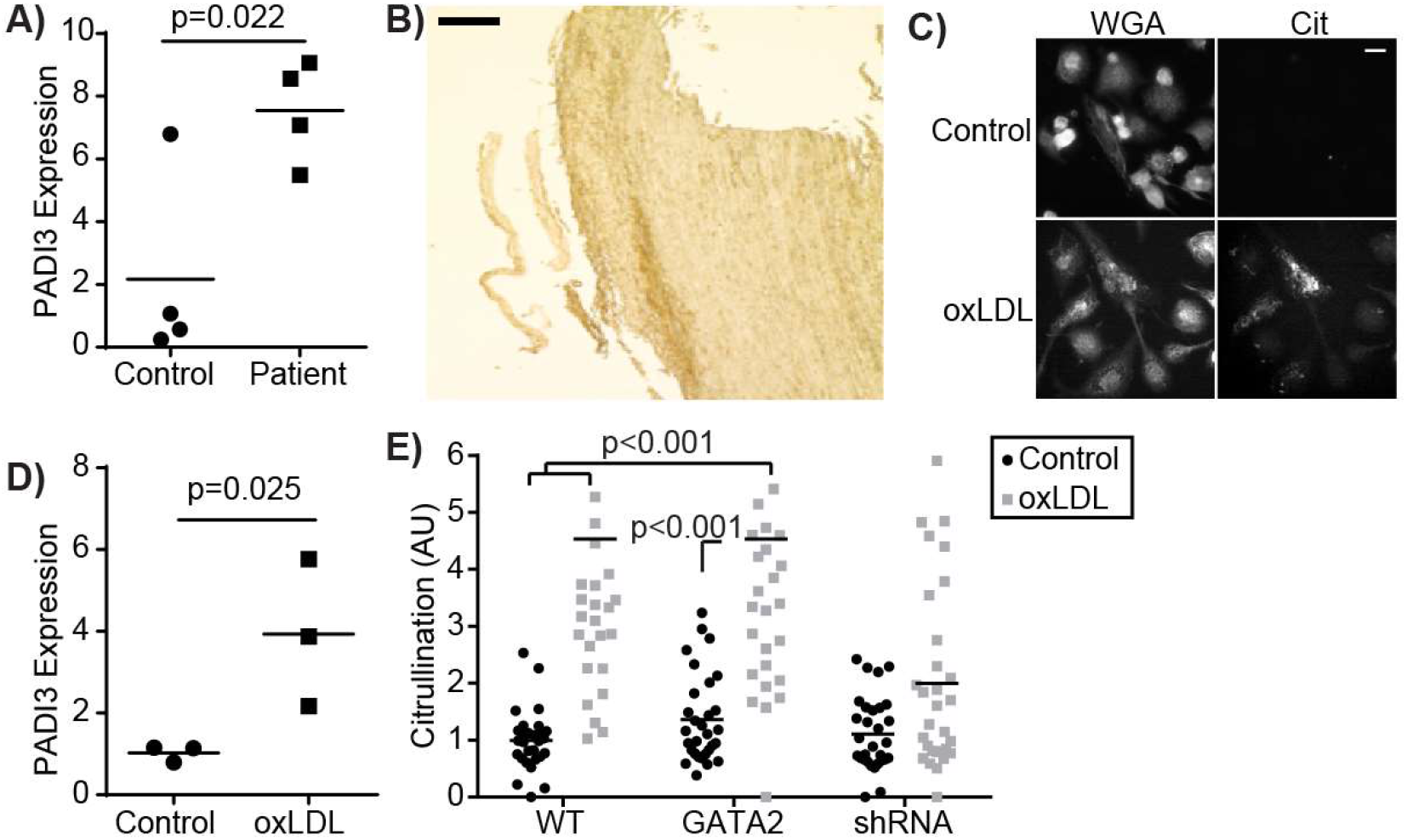
GATA2 Expression is Required for Plaque Citrullination. **A)** RT-PCR quantification of PADI3 expression in intima-derived macrophages (patient) versus age- and sex-matched monocyte-derived macrophage controls (Control). **B)** Low-magnification image of an aortic punch biopsies stained for citrullinated proteins. Scale bar is 1 mm. **C)** THP1 derived macrophages that are either normocholesterolemca (Control) or hypercholesterolemic (oxLDL) stained with a plasma-membrane marker (WGA) and an anti-citrulline antibody (Cit). Scale bar is 10 μm). **D)** RT-PCR quantification of PADI3 expression in THP1-derived macrophages cultured under normocholesterolemic (Control) or hyperchoelsterolemic (oxLDL) conditions. **E)** Protein citrullination in THP1-derived macrophages expressing a vector control (WT), GATA2 transgene (GATA2) or GATA2-targeting shRNA (shRNA) in response to normocholesterolemic (Control) or hyperchoelsterolemic (oxLDL) conditions. Data is representative of (A,C) or quantifies as the ensemble of 4 (A-B) or 3 (C-E) independent experiments. p values were calculated using a Students t-test (A,D) or ANOVA with Tukey correction (E).

## Discussion

In this study we have identified a previously unreported macrophage polarization state present in the early stages of human atheroma formation. This polarization state differs greatly from other previously reported atherosclerosis-associated polarization states, most of which have been identified in advanced plaque or in animal models of disease. The polarization state we have identified is consistent with macrophages transitioning from an atheroprotective to an atherogenic phenotype, with genes regulating atheroprotective processes including cholesterol homeostasis and efferocytosis downregulated in these cells, but with these cells lacking the upregulation of inflammatory genes observed in macrophages from more advanced stages of disease. This polarization state was driven, in part, by the upregulation of the transcription factor GATA2, which was responsible for the impairment of both efferocytosis and the subsequent degradation of the engulfed material. These data identify GATA2 as an important regulator of macrophage dysfunction in early human atherosclerosis and provides a detailed map of the transcriptional changes occurring in macrophages during the early stages of plaque formation.

Macrophages are phenotypically plastic cells that perform a broad range of homeostatic, immunological, and wound-healing activities. The broad functional capabilities of macrophages are created by flexible transcriptional programs driven by tissue-specific cues, inflammatory cues in the local microenvironment, age-associated transcriptional changes, and the developmental history of the macrophage^24,56^. These polarization states are also highly plastic and respond to changes in the local environment. For example, classically polarized M1 and M2 macrophages undergo rapid repolarization via a Nrf2-driven transcriptional program into an atherogenic polarization state upon exposure to moLDL^31^. In this study, we determined that macrophages isolated from early-stage human atheromas lack upregulation of classical M1 (CD80/86, iNOS) or M2 (Arg1, CD206) markers, or markers previously reported in macrophage polarization states present in late-stage plaque (TREM2, Nrf2, Atf1)^27,30,31^. Instead, these cells downregulated many genes involved in the processes of efferocytosis, cholesterol processing, and the degradative pathway required for proper removal of both apoptotic cells and moLDL, while upregulating immunogenic enzymes such as PADI3. This expression pattern is consistent with the events proposed to occur in early-stage plaque. Indeed, the earliest histologically definable stage of atherosclerosis is the appearance of lipid-laden macrophages within the vascular intima^33^ – consistent with the decreased expression of genes involved in cholesterol trafficking (NPC1/2), efflux (ABCA1) and intracellular cholesterol homeostasis (SOAT1, NCEH1) that we observed within intima-infiltrating macrophages. Similarly, these cells have decreased expression of many genes required for the uptake and processing of apoptotic cells, setting the stage for the accumulation of necrotic foam cells which typifies later stages of atheroma development^47,57–59^.

Unlike T cells, in which polarization tends to be driven by one or a small number of transcription factors, the polarization of macrophages is generally a product of multiple cooperative transcription factors^60^. Consistent with this, we identified 20 transcription factors which were upregulated in intima-infiltrating macrophages. Eight of these transcription factors had no known function, and five were associated with cell cycle – consistent with previous observations that lesion-resident macrophages undergo proliferation and self-renewal during atherogenesis^61^. This left seven transcription factors potentially driving macrophage polarization in early plaque (MSX2, GATA2, FOXD4L5, RARγ, HSFX1, NKX6-1 and CIC). Of these, GATA2 was of particular interest as single nucleotide polymorphisms in this gene have previously been associated with cardiovascular disease through genome-wide association studies, although the mechanism by which these SNPs contribute to disease remains unknown^39^. Our data suggests that some of these SNPs may increase disease risk through inducing or enhancing the expression of GATA2 in macrophages in the vascular intima. GATA2 is a member of the GATA transcription factor family and functions as both a transcriptional activator and repressor^62^. GATA2 normally mediates the early stages of myelopoiesis, and must be downregulated to allow for completion of monocyte development^40,41,63^. GATA2 is not typically expressed by mature myeloid cells, although it is expressed in endothelial cells where it is required for the development and stability of vascular and lymphatic structures^64–67^. The importance of GATA2 in myelopoiesis is illustrated by patients with inactivating mutations in GATA2. A number of these patients develop MonoMAC syndrome, a primary immunodeficiency characterized by a near absence of circulating myeloid, NK and B cells, and as a result are highly susceptible to mycobacterial and Epstein-Barr Virus infections^67,68^. Very little is known of GATA2’s role in myeloid cell activity in peripheral tissues or during immune responses, with the exception of one study demonstrating that GATA2 expression in alveolar macrophages regulates fungal phagocytosis^69^. Our data demonstrating that GATA2 is upregulated in response to oxLDL, that GATA2 overexpression enhances cholesterol efflux, and that GATA2 upregulation impairs efferocytosis and the processing of efferocytosed cargo, suggests that GATA2 may be upregulated in response to the cellular stress caused by cholesterol accumulation, but is ultimately maladaptive through disrupting other atheroprotective macrophage functions. While it is tempting to hypothesize that GATA2 expression may revert macrophages to an earlier developmental state, the transcriptional profile we have identified in these cells correlates poorly with the reported transcriptome of developing myeloid cells^70^.

We examined gene expression changes in patient macrophages by microarray using M0-polarized macrophages differentiated *in vitro* from peripheral blood monocytes as a control. The lack of a more comparable source of control macrophages is a significant drawback of this approach, yet a suitable source of control cells is difficult to identify. Macrophages undergo significant age-associated changes in their gene expression profile, necessitating age-matched controls^25^. While there is a population of macrophages in the healthy artery, these are primarily located within the adventitia, with only 2% of arterial macrophages in the intimal layer^22,71^. Moreover, these macrophages are of embryonic origin, whereas plaque-resident macrophages are predominantly hematopoietic in origin^22,72,73^. Thus, the use of monocyte-derived macrophages from healthy age- and sex-matched controls recapitulates the correct ontology and age but lacks the tissue-specific cues encountered in the vascular intima, and our results must be interpreted accordingly. Another drawback of our approach is that while the plaques we identified in our aortic punch samples appear to be in the early stage of development from a histological perspective, they are isolated from patients with advanced disease in the coronary circulation. Ergo, some of the observed gene expression changes may be due to systemic effects of atherosclerosis, rather than changes specific to early-stage plaque development. Nonetheless, we observed differential expression of over 3,000 protein-coding genes in intima-infiltrating macrophages compared to controls. Undoubtedly, some of this difference is due to the nature of our controls, however, many of the features that we identify as unique in the pre-atherosclerotic, lesion-resident macrophage population are consistent with our expectations of macrophage dysfunction during early stages of atherosclerotic disease, and are not obviously connected to inherent differences between tissue versus cultured macrophages^34–37,60,74^. These features include marked downregulation of genes involved in cholesterol homeostasis, phagocytosis/efferocytosis, phagosome/efferosome maturation, and antigen processing and presentation. Rather, we contend that these features define a novel macrophage phenotype associated with early, pre-atherosclerotic lesions.

In this study we are the first to report that GATA2 is upregulated in macrophages in early-stage human atherosclerotic plaque and is responsible for inducing both efferocytic defects and autoantigen-generating processes in these cells that contribute to the development of atherosclerosis. This GATA2-mediated dysregulation of macrophage activity is part of an aberrant polarization state which arises early in plaque development and sets the stage for disease progression. As such, targeting of GATA2 and the processes downstream of GATA2 activation may represent a novel therapeutic opportunity to limit plaque development by targeting defective efferocytosis within the atherosclerotic plaque.

## Materials and Methods

### Materials

pBabePuro-GATA2 (Addgene #1285) was a gift from Gokhan Hotamisligil^75^. pLXV-zsGreen lentiviral vector and pGFP-C-shLenti vector were gifts from Dr. Jimmy Dikeakos. Coverslips, slides, Leiden chambers and 16% paraformaldehyde (PFA) were from Electron Microscopy Sciences (Hatfield, PA). DMEM, RPMI and fetal bovine serum (FBS) were from Wisent (Montreal, Canada). Restriction enzymes, Gibson assembly reagent, and ligase was from New England Biolabs (Whitby, Canada). M-CSF, GM-CSF, INFγ and IL-4 were purchased from Peprotech (Montreal, Canada). Micro-beads were from Bangs Laboratories (Fishers, Indiana). Lipids and cholesterol were from Avanti polar lipids (Alabaster, AL). Lympholyte-poly and all secondary antibodies/Fab’s were from Cedarlane Labs (Burlington, Canada). pHrodo, NBT, human oxLDL, TRITC-Dextran, nigericin, Hoechst 33342, NanoDrop 1000 Spectrophotometer, Fast SYBR Green Master Mix, QuantStudio 5 Real-Time PCR System and the Anti-Citrulline (Modified) Detection kit were from ThermoFisher Canada (Mississauga, ON). Human IgG, PMA, anti-modified citrulline antibody were from Sigma-Aldrich (Oakville, ON). Anti-CD163, anti-GATA2 and anti-CD68 were from Abcam (Toronto, Canada). 4.0 mm CleanCut RCL Aortic Punch was from QUEST Medical (Allen, Texas). Gemini Fluorescence Microplate Reader running SoftMax Pro was from Molecular Devices (San Jose, California). PureZOL RNA isolation reagent and iScript Select cDNA Synthesis Kit were from BioRad (Hercules, California). CapSure HS LCM Caps and GeneChip WT Pico Reagent Kit were from Applied Biosystems (Forest City, California) Prism 6 software was from GraphPad (La Jolla, California). FIJI was downloaded from https://fiji.sc/ ^76^. All other materials were purchased from Bioshop Canada (Burlington, Canada).

### Human Plaque Histology & Macrophage Isolation

Patient tissues used in this study were obtained under a discarded tissue protocol from patients undergoing elective coronary artery bypass graft surgery at London Health Sciences Centre, London, Canada. This study was reviewed and approved by the Office of Human Research Ethics at Western University Health Sciences Research Ethics Board (HSREB Reference Number: 107566) and all procedures were performed in accordance with the guidelines of the Tri-Council policy statement on human research. Since this is a discarded tissue study, we did not collect any personal health information, personal identifying information or clinical data other than the age range and the male-to-female ratio of the entire cohort. Researchers were blinded to patient identification and clinical characteristics. Aortic punch tissue specimens were obtained intra-operatively by a cardiac surgeon using a 4.0-mm diameter aortic punch and placed in cold saline. Specimens were then bisected evenly using clean surgical scissors, with one half of the tissue used to prepare frozen sections and the other for preparing paraffin sections. For laser capture microdissection, tissue specimens were embedded in OCT freezing compound and frozen on dry ice over a period of 5-10 min within 30 min of collection, then placed into −80 °C for storage. For immunofluorescence and Oil-Red-O (ORO) staining, tissue specimens were fixed in fresh 4% paraformaldehyde (PFA) for 24 hr at 4 °C, placed in PBS + 15% sucrose until the tissue lost buoyancy (~4 hr), and then placed into PBS + 30% sucrose overnight at 4 °C. For all other stains, paraffin-embedded sections were fixed 4% PFA for 24 hr at 4 °C, then stored in 70% ethanol. Specimens were dehydrated by 1 hr immersion in 70% and 95% ethanol, followed by four immersions in 100% ethanol (1, 1.5, 1.5 and 2 hrs). Sections were cleared with two 1 hr immersions in xylene, then immersed two times 1 hr in paraffin wax (58 °C). Processed tissues were embedded into paraffin blocks.

OCT-embedded tissues were sectioned into 10 μm sections and paraffin-embedded samples at 5 μm, and placed onto clean, RNAse-free slides. All histology (H&E, ORO, Movat’s pentachrome stain and TUNEL stain) was performed at the Robarts Molecular Pathology core.

For laser capture microdissection (LCM), slides were stored at −80C until processing. Slides were fixed in ice-cold acetone for 2 min, air-dried for 30 sec and stained with primary antibody (anti-CD163, 30-60 μg/mL, 3 min), washed 2× with PBS, and stained with a secondary antibody (1:100 dilution, 3 min). CD163 was used in lieu of CD68, as CD163 is found on all human macrophages where as CD68 stains cells in the atheroma of both macrophage and smooth muscle origin^38^. After two PBS washes, the slides were dehydrated by sequential addition of 75%, 95% and 100% ethanol (30 sec/step) and dried by immersion in xylene for 5 min. LCM was performed on an ArcturusXT Laser Capture Microdissection System (Applied Biosystems) using CapSure HS LCM Caps. Samples were captured using IR capture laser of 15 μm diameter at 70 mW, 1,500 μsec and 1 hit per capture. A minimum of 200 captures per section across four consecutive sections was performed to obtain sufficient cell numbers for downstream analysis.

### Microscopy & Image Analysis

Unless otherwise noted, all microscopy was performed using a Leica DMI6000B microscope equipped with 40×/1.40NA, 63×/1.40NA and 100×/1.40 NA objectives, photometrics Evolve-512 delta EM-CCD camera, heated/CO_2_ perfused stage, Chroma Sedat Quad filter set with blue (Ex: 380/30, Em: 455/50), green (Ex: 490/20, Em: 525/36), red (Ex: 555/25, Em: 605/52) and far-red (Ex: 645/30, Em: 705/72) filter pairs, and the LAS-X software platform. All image analysis was performed in FIJI^76^. For quantitative imaging, the same exposure intensity, exposure time and EM-CCD gain setters were used within a single experiment, and control-normalized data used for comparisons between experiments.

### Primary Macrophage Culture

The collection of blood from healthy donors was approved by the Health Science Research Ethics Board of the University of Western Ontario and was performed in accordance with the guidelines of the Tri-Council policy statement on human research. Blood was drawn from age- and sex-matched volunteers without diagnosed coronary artery disease by venipuncture. Blood was collected with heparinized tubes, 5 ml of blood layered over 5 ml of Lympholyte-poly, and centrifuged at 300 × g for 35 min. The top band of cells was collected, washed once (300 × g, 6 min, 20 °C) with phosphate buffered saline (PBS, 137 mM NaCl, 10 mM Na_2_HPO_4_), and resuspended at 2 × 10^6^ cells/ml in RPMI-1640 + 10% FBS + 1% antibiotic–antimycotic. 200 μl of this suspension was placed on sterile glass coverslips for 1 h at 37 °C, washed twice with PBS, and then differentiated into M0-polarized macrophages as per our published protocols^52,77^.

### Microarray

Total RNA was prepared from patient aortic punch macrophages or M0-polarized control macrophages by TRIzol extraction. RNA quality was checked using a 2100 Bioanalyzer Instrument (Agilent, Santa Clara, California), with samples below an A260/280 of 1.5 and/or RIN of 7 rejected from analysis. A minimum of 2.0 μg of total RNA was from each sample was used in the microarray. RNA samples were prepared for array hybridization using the GeneChip WT Pico Reagent Kit according to the manufacturer’s instructions with 12 cycles of pre-*in vitro* transcription amplification. Poly-A RNA standards provided by the manufacturer were used as exogenous positive controls. Samples were analyzed on the GeneChip Human Gene 2.0 ST Array chip hybridized on the GeneChip Scanner 3000 7G System (Applied Biosystems) according to the manufacturer’s recommended instructions. Raw microarray data were analyzed using the Partek Genomics Suite platform (Partek), with reads normalized by a Robust Multi-array Average (RMA) procedure with quantile normalization. Mixed-model ANOVA was used to identify differentially expressed genes with a foldchange cutoff of 2.0 and p-value cutoff of p<0.05. Unsupervised hierarchical clustering of all differentially expressed genes was performed to generate a heat map of differential gene expression. Results were verified using Bioconductor^78^. The oligo package was used to read in raw microarray data files and perform RMA normalization, and the limma package used to identify differentially expressed genes using a linear model approach along with an empirical Bayes method to better estimate errors in log-fold change.

Partek Genomics Suite was used to perform gene ontology microarray data using GO ANOVA, with analysis restricted to groups with more than two and fewer than 150 genes. The top 100 GO biological function terms were visualized using the REVIGO online software platform (Rudjer Boskovic Institute, Zagreb, Croatia). Gene set enrichment analysis (GSEA) was performed using the Gene Set Enrichment Analysis software platform (Broad Institute, Cambridge, Massachusetts) with publicly-available annotated gene sets (cholesterol_homeostasis, antigen_processing_presentation, fcr_mediated_phagocytosis, phagosome_maturation) obtained from the Molecular Signatures Database (MSigDB). Analysis was run using 10,000 permutations with FDR < 0.25.

### THP-1 Cell Culture

THP-1 human monocytes were cultured in DMEM + 10% FBS, and split upon reaching a density of ~2 × 10^6^/ml. To generate THP-1 derived macrophages, #1.5 thickness, 18 mm diameter circular coverslips were placed into the wells of a 12-well plate, 1 × 10^5^ THP-1 cells placed into each well cultured for 72 hrs in DMEM + 10% FBS + 100 nM PMA. Cells were then incubated with human oxLDL at a concentration of 100 μg/mL for 72 hrs (THP-1 cells^79^). oxLDL-treated cells were washed with PBS to remove excess oxLDL and were placed into fresh, oxLDL-free media prior to use.

### Cloning

GATA2 was sub-cloned from pBabePuro-GATA2 into the pLXV-zsGreen lentiviral vector by restriction cloning, using EcoRI as per the manufacturer’s instructions. GATA2 shRNA was ordered as an oligo (5’-AAGGA TCCAG CAAGG CTCGT TCCTG TTCAT CAAGA GTGAA CAGGA ACGAG CCTTG CTTTT TTACC GGTAA-3’) and cloned into into pGFP-C-shLenti by restriction cloning with BamHI and AgeI. These were packaged into a VSV-G pseudotyped lentiviral vector using HEK 293T cells expressing pMD2.G (2 ng, Addgene #12259), pCMV-DR8.2 (5 ng, Addgene #12263) and 2.5 μg of the lentiviral vector. 72 hrs post transfection, virus was purified and stored as described previously −80°C ^80^. THP-1 cells were transduced by adding 500 μL of viral isolate to 1 × 10^6^ of THP-1 cells with 8 μg/mL polybrene. One week later, transduced cells were fluorescence-activated cell-sorted based on their expression of the zsGreen/GFP marker encoded by the vectors. All primer and shRNA sequences can be found in Supplemental Table 2.

### Immunoblotting

At least 5 × 10^5^ cells were lysed in Laemmli’s buffer + 10% β-mercaptoethanol and Halt protease inhibitor cocktail, boiled briefly, separated on a 10% SDS-PAGE gel, and transferred to nitrocellulose membrane. The membrane was blocked for at least one hour with Trisbuffered saline with 0.1% Tween 20 (TBS-T) + 5% bovine serum albumin in PBS, incubated for 1 hr with anti-GATA2 antibody (1:500 dilution) or anti-GAPDH (1:1000) in TBS-T + 2.5% milk powder. The blot was washed 3 × 15 min in TBS-T, and then a 1:10,000 dilution of an appropriate IR700 or IR800 secondary antibody added for 30 min to 1 hr in TBS-T + 5% bovine serum albumin in PBS. The blots were washed 3 × 15 min washes in TBS-T and imaged using an Odyssey CLx (LI-COR Biosciences, Lincoln, Nebraska).

### Real-Time PCR

Total RNA was isolated from cells or patient tissues through acid guanidinium thiocyanate-phenol-chloroform extraction using PureZOL RNA isolation reagent. Cells or tissues were suspended in PureZOL, vortexed briefly, and incubated for 5 min at room temperature. Chloroform was added 1:5 v/v and incubated for another 5 min with intermittent agitation. Samples were centrifuged for 15 min at 12,000 ×g and 4 °C, then the aqueous layer transferred to a fresh tube to which an equal volume of isopropanol and 0.5 μL of 20 mg/mL glycogen were added. Samples were incubated at −80 °C for at least 30 min, centrifuged for 15 min at 12,000 ×g and 4 °C, and the RNA pellet washed 2× with 75% ethanol, air dried for approximately 5 min at room temperature and reconstituted in a minimal amount of RNase-free ddH_2_O. RNA concentration and quality were measured using a NanoDrop 1000 Spectrophotometer. cDNA was generated using the iScript Select cDNA Synthesis Kit according to the manufacturer’s instructions, with an equal amount of starting RNA an equal mix of random and oligo(dT)20 primers. cDNA concentration and quality were checked using a spectrophotometer prior to use in qPCR reactions. qPCR was performed using the Fast SYBR Green Master Mix and an equal amount of starting cDNA. PCR reactions were run on a QuantStudio 5 Real-Time PCR System for 40 cycles. Relative transcript express was calculated using the ΔΔCt method with 18S used as the housekeeping gene. All RT-PCR primer sequences can be found in Supplemental Table 2.

### Phagocytosis and Efferocytosis Assays

Phagocytosis and efferocytosis assays were performed as described previously^77,81^. Briefly, phagocytic targets were generated by washing 10 μl of 5 μm diameter P(S/DVB) polystyrene beads (6,000×g/1 min) with 1 ml of PBS. Beads were suspended in 100 μl of PBS + 0.1 mg/ml human IgG and incubated at 20°C for 30 min. Beads were then washed as above and suspended in 1 ml of DMEM. Efferocytic targets were prepared by suspending a 4 μmol mixture of 20% phosphatidylserine, 79.9% phosphatidylcholine and 0.1% biotin-phosphatidylethanolamine in chloroform, and to this adding 10 μl of 3 μm diameter silica beads. After a 1 min vortex, the chloroform was evaporated with nitrogen and the beads suspended in 1 ml of PBS. The beads were then washed 3× as described above, and then suspended in 1 ml of DMEM. These beads were then added to macrophages at a 10:1 bead:macrophage ratio, spun at 200 × g for 1 min to force contact between the cells and beads, and then incubated for the indicated time. The samples were then fixed with 4% PFA for 20 min, washed, and if required, non-internalized beads detected by incubation for 20 min in PBS + 1:1000 dilution of anti-human Cy3secondary antibody, or DMEM + 1:500 dilution of Alexa-555 labeled streptavidin. Samples were then washed 3x in PBS or DMEM and mounted on slides with Permafluor and imaged.

### Lysosome Fusion

Lysosomes in THP-1 derived macrophages were loaded with 100 μg/mL 10,000 MW TRITC-conjugated dextran for 16 hrs, followed by a 90 min chase with serum-free RPMI. IgG-coated beads, prepared as above, were added at a 10:1 bead:macrophage ratio and briefly centrifuged for 1 min at 400 ×g, then incubated for 30 min at 37 °C and 5% CO_2_. Following incubation, cells were washed 1× with PBS and labelled 1:1,000 with a Cy5-conjugated anti-human IgG secondary antibody to label noninternalized beads. Cells were washed 3× 5 min with PBS, fixed with 4% PFA for 20 min, washed an additional 3× 5min and mounted onto glass slides using Permafluor. Samples were imaged at 63× magnification, using the red (dextran) and far-red (beads) channels.

### Phagosome pH

Human IgG was labeled with pHrodo as per the manufacturer’s instructions, and phagocytic targets prepared as described above using a 500:1 ratio of pHrodo-IgG:Alexa-647-labeled irrelevant antibody. Live cell microscopy was performed at 63× magnification, using point-visiting to image 4-5 locations (20-30 cells) per condition. The media was then replaced with DMEM + phagocytic targets and a 90 min timelapse captured at 2 min/frame using the DIC, red (pHrodo) and far-red filter sets, maintaining the same camera and exposure settings used across all experimental conditions within an individual experiment. At the end of the experiment the media was replaced with high-potassium media + 10μg/ml nigericin at a pH pf 4.0, 5.0, 6.0 and 7.0. The cells were imaged after each media change, thereby providing an *in situ* pH calibration for each phagosome. The resulting images were imported into FIJI, individual beads tracked using the manual tracking plugin and the pHrodo:Alexa-647 ratio measured in each phagosome at each time point and the pH determined using the data from the calibration images. Phagosomal FITC staining was normalized to the integrated FITC intensity across the whole cell at the first time point.

### Oxidant Production

A fluorescence microscopy-based NBT assay protocol was used to assess macrophage NADPH oxidase activity^82^. Apoptotic Jurkat cells were generated as per our published protocls^77,81^ and added to PMA-differentiated THP-1 cells at a 10:1 Jurkat:macrophage ratio. Samples were centrifuged for 1 min at 400 ×g to force contact between the macrophages and ACs, and NBT was added to the medium to a final concentration of100 μg/mL. After 1 hr incubation at 37 °C and 5% CO_2_ cells were washed 3× 5 min and fixed with 4% PFA for 20 min. Cells were washed 3× 5 min with PBS and mounted onto slides using Permafluor. Fluorescence of diformazan deposits formed were imaged on using at 63×, using the far-red fluorescence channel.

### Cholesterol Uptake

Cholesterol accumulation was measured by staining with either ORO or Nile Red (NR). ORO solution (1:250 w/v ORO powder in isopropanol) was diluted 3:2 in ddH_2_O and passed through a 0.2 μm filter. Cells were washed 1× with PBS, fixed with 4% PFA for 20 min, washed an additional 3×, washed briefly with 3:2 v/v mixture of isopropanol and ddH_2_O, and allowed to dry. Cells were then covered with the minimum volume of ORO working solution and allowed incubated for 5 min, washed 3× 5 min with PBS prior to mounting for imaging. ORO accumulation was quantified by extracting stained cells with 100% isopropanol for 10 min at room temperature with gentle agitation, and absorbance measured at 518 nm. NR (1 mg/ml in acetone) was diluted 1:100 into PBS or serum-free DMEM. For fixed cell staining, the PBS solution was added to fixed and washed cells for 5 min, followed by 3× 5 min with PBS. For live-cell staining, the DMEM solution was added to cells in a Leiden chamber, and images acquired every 15 min for 8 hrs at 63× magnification, using the red fluorescence channel.

### Cholesterol Efflux

Cholesterol efflux was measured as described by Sankaranarayanan *et al*^83^. Briefly, BODIPY-cholesterol and unlabeled cholesterol (1:5 v/v) were mixed and dried under nitrogen, solubilized in MEM-HEPES (MEM media with 10 mM HEPES, pH 7.4) + 20 mM methyl-β-cyclodextran (MβCD)) at a molar ratio of 1:40 cholesterol:MβCD. The mixture sonicated at 37 °C for 30 min, placed onto a stirring hot plate pre-at 37 °C for 3 hrs. Immediately prior to use this labeling media is sonicated at 37 °C for 30 min and filtered through a 0.45 μm filter. Apolipoprotein B (ApoB)-depleted human serum was prepared by adding 1:40 v/v 1M CaCl_2_ to whole human plasma, incubated for 1 hr, and serum separated by centrifugation for 5 min at 1,000 ×g. ApoB was then precipitated by addition of 20% PEG 8,000 in 200 mM glycine buffer, pH 7.4, at a 2:5 v/v ratio. The mixture is allowed to incubate for 20 min and then centrifuged for 30 min at 10,000 ×g and 4 °C.

THP-1 cells were cultured in a 48-well plate to a density of 75,000 cells/well and differentiated into macrophages by culture with 100 nM PMA for 24 hrs prior. Cells were washed 1× with PBS and incubated with 250 μL/well of labelling media for 1 hr, washed 2× with MEM-HEPES, and then incubated with serum-free RPMI + 0.2% BSA for 16 hrs. Cells were washed 2× with MEM-HEPES, then incubated with MEM-HEPES + 10% apoB-depleted serum for 4 hrs. All procedures were performed at 37 °C and 5% CO_2_. Media from each well was collected, filtered through a 0.45 μm filter and fluorescence intensity (ex. 482 nm, em. 515 nm) was recorded using a Gemini Fluorescence Microplate Reader (Molecular Devices). Negative controls were solubilized with 4% cholic acid for 4 hrs, the supernatants filtered through a 0.45 μm filter, and fluorescence intensity was recorded and used as the baseline value for total cholesterol present within the cells. The percent cholesterol efflux was calculated as fluorescence intensity of media divided by fluorescence intensity of cholic acid.

### Citrullination

Cells or tissues sections were incubated with wheat germ agglutinin (1:200 in PBS), 10 min at room temperature, then washed three times in PBS and permeabilized for 10 min with a 0.2% Triton X-100 solution at room temperature. Cells were then washed three times in PBS, incubated with equal volumes of modification reagent A (0.5% FeCl2, 4.6 M sulfuric acid, and 3.0 M phosphoric acid) and modification reagent B (0.5% 2,3-butanedione monoxime, 0.25% antipyrine, 0.5M acetic acid) at 37 °C for 3 h. Cells were then washed three times in PBS, blocked for 1 hr with 5% FBS, then incubated for 1 h with anti-modified citrulline or an isotype control at 1 μg/ml in 5% FBS. Cells were washed three times in PBS and incubated with a 1:1000 dilution of Cy3-labeled anti-human secondary antibody in 5% FBS for 1 hr. Cells were washed three times with PBS, counter-stained with 2 μg/ml Hoechst for 10 min, washed a final three times and mounted on coverslips with Permafluor mounting medium.

### Statistics and Data Analysis

All datasets were tested for normal distribution using a Shapiro-Wilk test. Normally distributed data is presented as mean ± SEM and analyzed using a Students *T*-test or ANOVA with Tukey correction. Non-parametric data is presented as median ± interquartile range and analyzed using a Kruskal-Wallis test with Dunn correction. All statistical analyses were performed using Graphpad Prism 6.

## Acknowledgements

The authors would like to thank Dr. Edward J. Tweedie (Department of Pathology and Laboratory Medicine, The University of Western Ontario, London, Ontario, Canada) for his assistance with the histological classification of the aortic biopsy samples. David Carter (London Regional Genomics Centre, Robarts Research Institute, London, Ontario, Canada) for his assistance with the microarray experiments and data analysis, Caroline O’Neil (Molecular Pathology Core Facility, Robarts Research Institute) for her help with tissue histology and laser capture microdissection, and Dr. Kristin Chadwick (London Regional Flow Cytometry Facility, Robarts Research Institute) for her assistance with FACS-based cell sorting. This study was funded by a Canadian Institutes of Health Research (CIHR) Operating Grant MOP-123419, an Ontario Ministry of Research and Innovation Early Research Award, and a University of Western Ontario Accelerator grant to BH. CY was funded by a Vanier PhD Scholarship and CIHR Md/PhD studentship. JDD was funded by a Canadian Institutes of Health Research Operating Grant MOP-389413. ENP was funded by an Alexander Graham Bell Doctoral Canada Graduate Scholarship from the Natural Sciences and Engineering Council. JA was funded by the Bone and Joint Institute at the University of Western Ontario. The funding agencies had no role in study design, data collection and analysis, decision to publish, or preparation of the manuscript.

