## Supplemental Figures and Tables for "GATA2 Expression by Intima-Infiltrating Macrophages Drives Early Atheroma Formation"

### Supplemental Materials

Charles Yin, Angela M. Vrieze, James Akingbasote, Emily N. Pawlak, Rajesh Abraham Jacob, Jonathan Hu, Neha Sharma, Jimmy D. Dikeakos, Lillian Barra, A. Dave Nagpal, Bryan Heit.

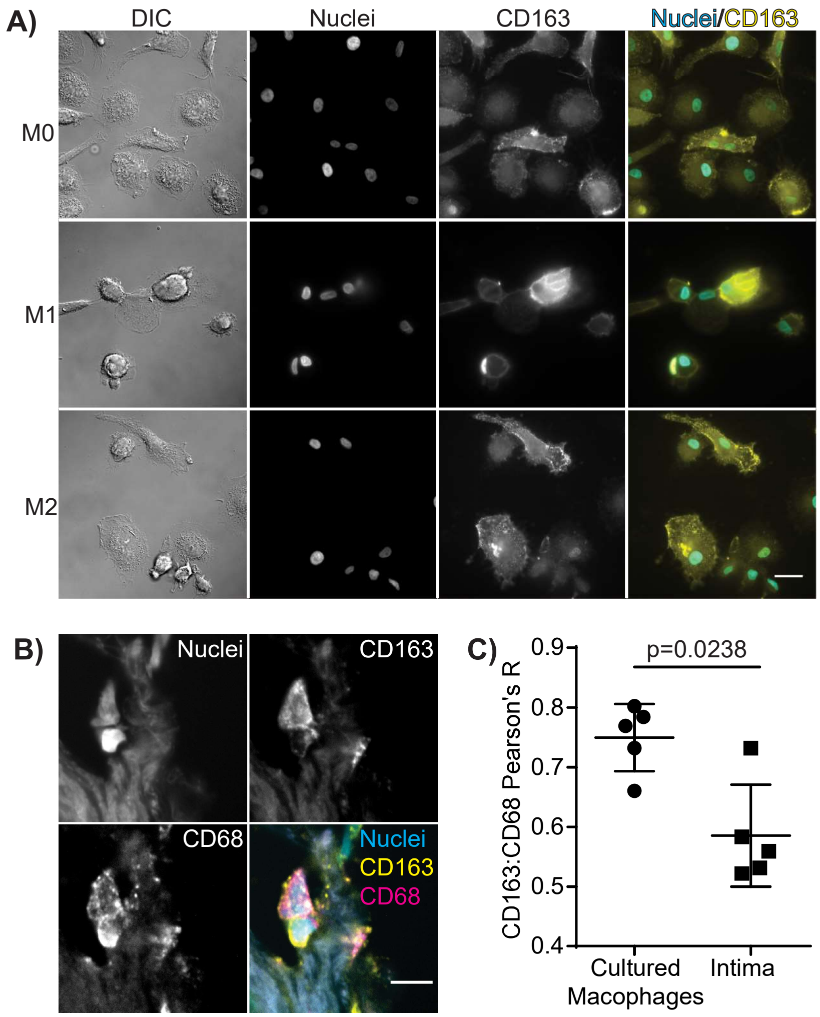


**Figure S1: Validation of CD163 as a Human Macrophage Marker. A)** Monocyte-derived macrophages were polarized *in vitro* to a M0, M1 or M2 polarization state and stained for nuclei with Hoechst and for CD163. Scales bars are 20 μm. **B)** Hoechst (Nuclei, cyan), CD163 (yellow) and CD68 (magenta) staining in frozen sections from human aortic punch biopsies confirmed to contain early-stage atherosclerotic plaque by Oil-Red-O and H&E immunohistochemistry of serial slices (not shown). Scale bar is 10 μm. **C)** Pearson's colocalization coefficient of CD163 with the canonical macrophage marker CD68 in cultured macrophages and in intima-infiltrating macrophages. Data is representative of, or quantifies, 5 independent experiments. p value was calculated using a Mann-Whitney test.

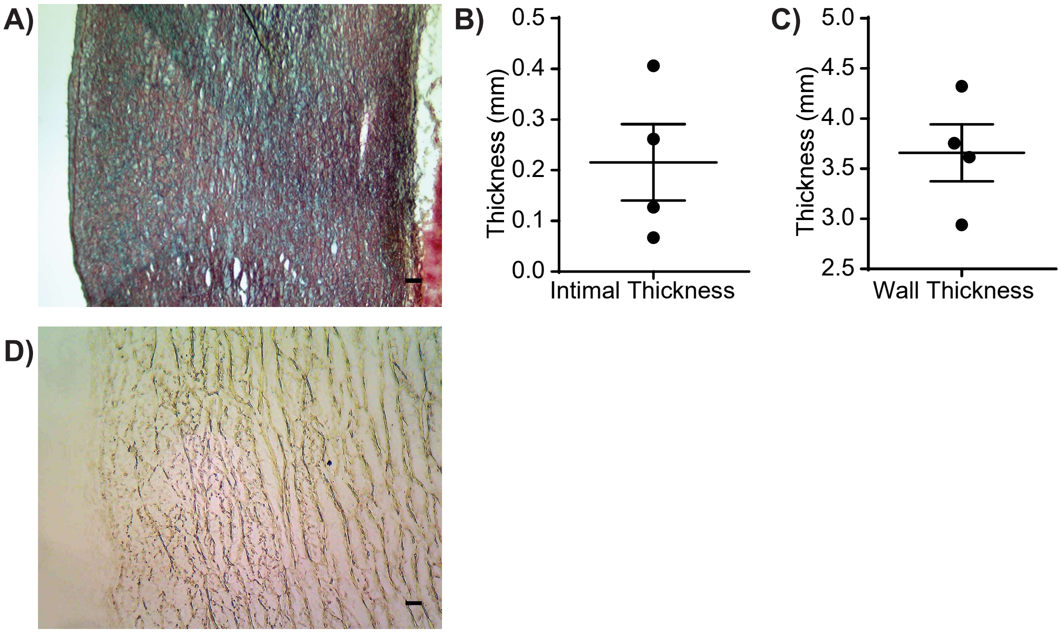


**Figure S2: Additional Histological Characterization of Aortic Punch Biopsies.** Human aortic punch biopsies were recovered from patients undergoing routine cardiac bypass graft surgery and cut into thin cryosections for histological analysis. **A)** Representative low-power field Movats stain, used to assess intimal **(B)** and wall **(C)** thickness. **D)** Representative TUNEL stain. Data is representative of, or quantifies, samples from 4 patients, Scale bars are 100 nm.

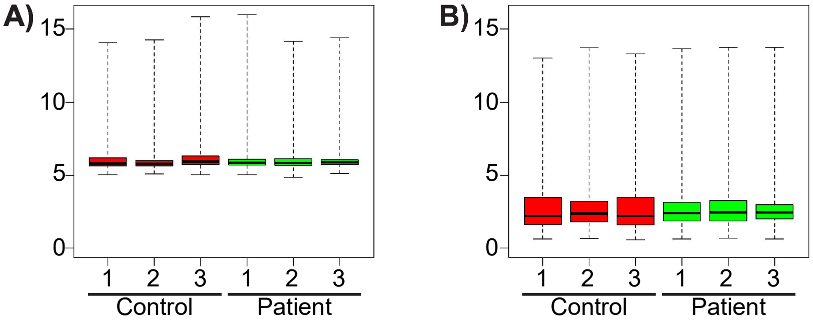


**Figure S3: Normalization Microarray Data. A)** Pre-normalization. **(B)** Post-normalization. Samples from 3 patients or 3 controls. Data is plotted as normalized microarray read intensities.

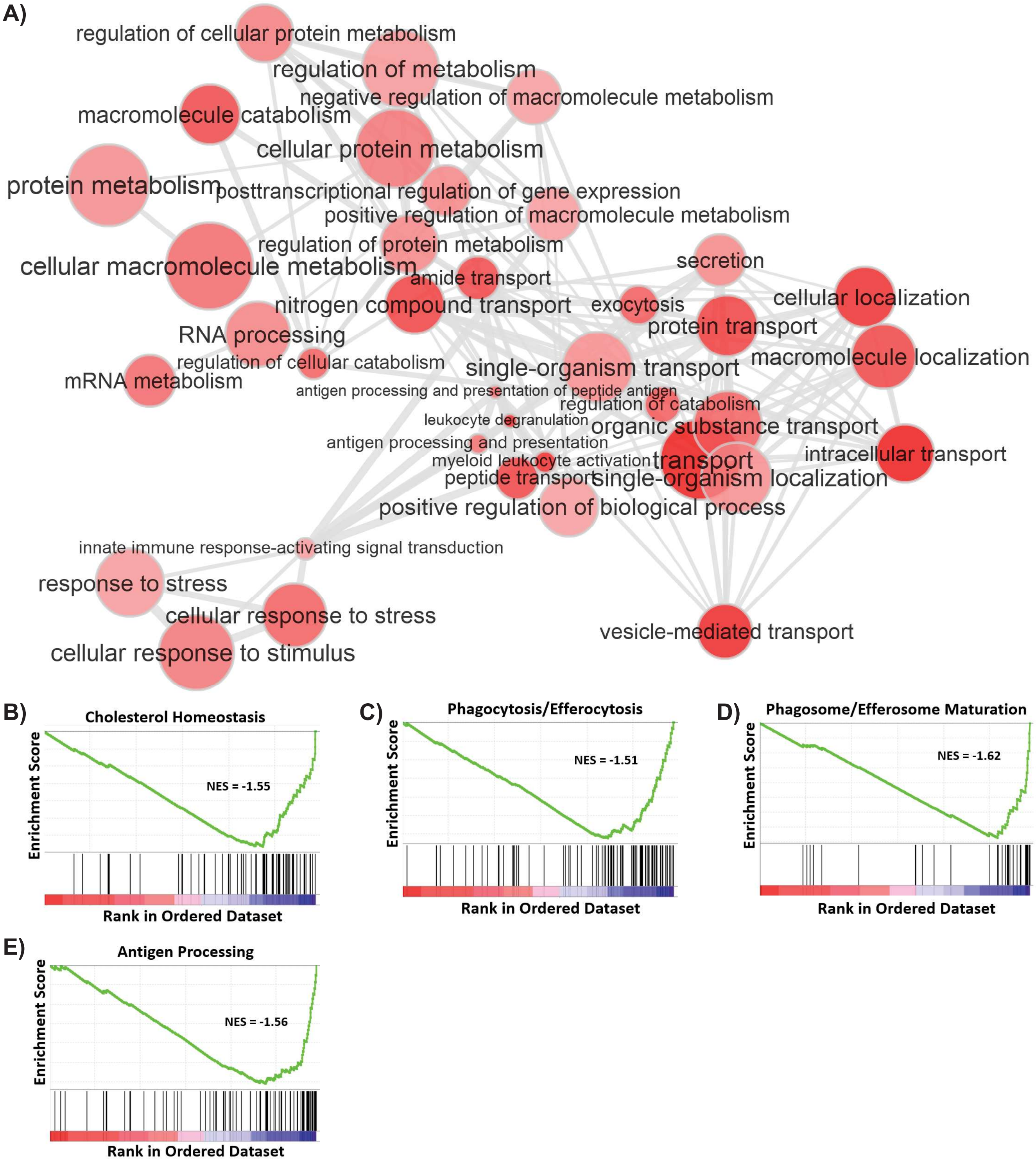


**Figure S4: Gene ontology and pathway enrichment analysis of intima-infiltrating macrophage gene expression patterns.** **A)** REViGO gene ontology analysis of intima-infiltrating macrophages reveals three major clusters of gene dysregulation: (top) catabolic processes including those regulating cholesterol homeostasis, (bottom left) pathways regulating responses to cell stress, and (bottom-right) the signaling and vesicular trafficking pathways regulating phagocytosis, endocytosis and efferocytosis. **B-E)** Pathway enrichment analysis identified the statically significant enrichment of genes involved in cholesterol homeostasis (B), the engulfment of pathogens or apoptotic cells (C), the degradation of phagocytosed or efferocytosed targets (D), and the subsequent generation and presentation of efferosome/phagosome-derived antigens (E). Data is calculated from the aggregated microarray data from 3 patients and 3 age- and sex-matched controls.

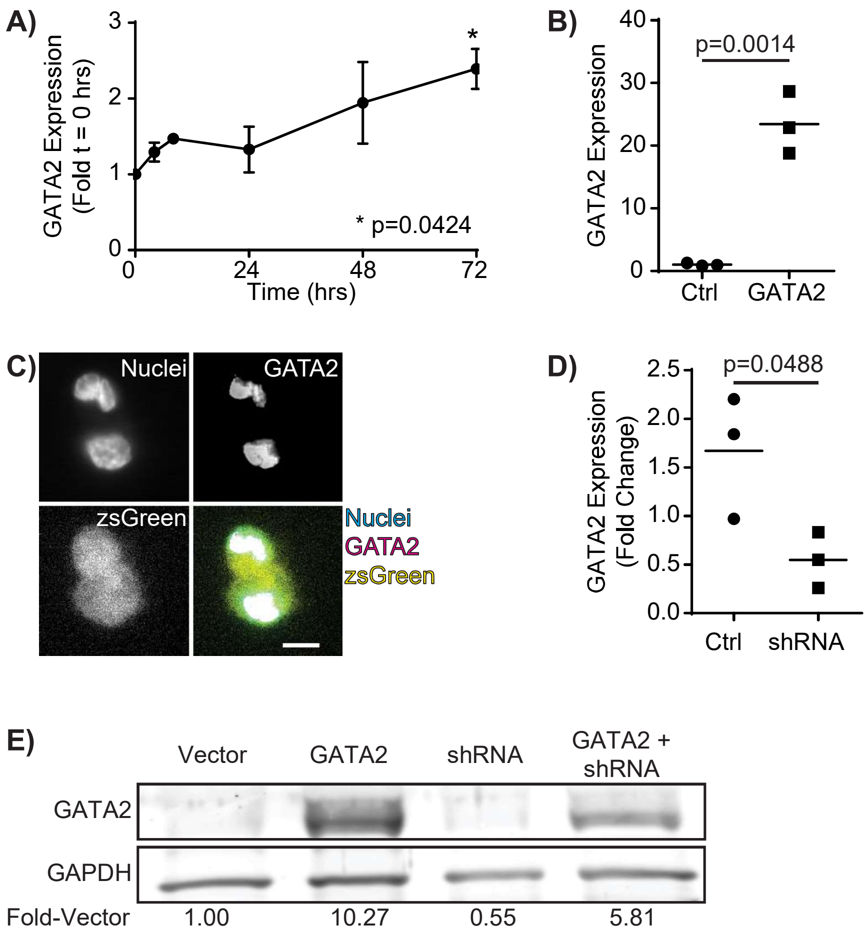


**Figure S5: Characterization of an *in vitro* THP1-Derived Macrophage hypercholesterolemia Model. A)** RT-PCR quantification of GATA2 expression over time in 100 μg/mL oxLDL treated THP1 derived macrophages. **B)** GATA2 expression in THP1-derived macrophages, as quantified by RT-PCR, transformed with either empty (Ctrl) or GATA2 lentiviral expression vectors. **C)** GATA2 immunostaining in THP1-derived macrophages expressing GATA2 from a lentiviral expression vector with a zsGreen selectable marker. **D)** Fold-change in GATA2 expression in THP1 macrophages expressing either a scrambled (Ctrl) or GATA2-specific shRNA following treatment with 100 μg/mL oxLDL for 72 hrs. Quantification is by RT-PCR. **E)** Immunoblotting of GATA2 protein in THP1-derived macrophages transduced with empty (Vector), GATA2-expressing (GATA2) or GATA2-shRNA expressing (shRNA) vectors. Fold-change in GATA2 protein levels, normalized to GAPDH expression and the vector control, are shown below each lane. Data are representative of (C,E), or quantify (A,B,D), 3 independent experiments. p values were determined using an ANOVA with Tukey post-hock test (compared to 0 hrs, A) or Students t-test (B,D).

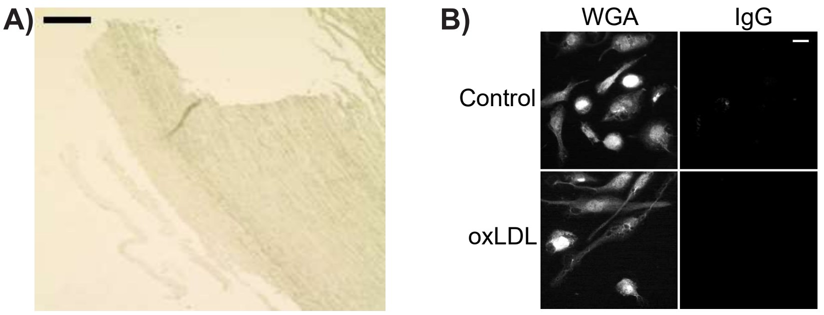


**Figure S6: Citrullination Staining Controls.** Serial section from the same aortic punch biopsy as Figure 6B **(A)**, or THP1-derived macrophages cultured under the same conditions as in Figure 6C-E **(B)** were stained using the same staining protocol as for anti-citrulline, but using an irrelevant IgG in place of anti-citrulline. Data is representative of 3 independent experiments. Scale bars are 10 mm (A) or 10 μm (B).

**Table 1:** Due to the large size of this table it has not been included in this preprint.

**Table S2: Primer and shRNA sequences**

| Gene | Sequence |
|---|---|
| | *qPCR primers* |
| 18S | 5'-GAGGGAGCCTGAGAAACGG-3'<br>5'-GTCGGGAGTGGGTAATTTGC-3' |
| CD14 | 5'-AGCCAAGGCAGTTTGAGTCC-3'<br>5'-TAAAGGACTGCCAGCCAAGC-3' |
| SMA | 5'-CCGACCGAATGCAGAAGGA-3'<br>5'-ACAGAGTATTTGCGCTCCG-3' |
| GATA2 | 5'-GTCACTGACGGAGAGCATGA-3'<br>5'-GGCACATAGGAGGGGTAGGT-3' |
| E4F1 | 5'-ACACCACACAGGCGAGA-3'<br>5'-TCCTCAGACACCAGCAAC-3' |
| RARG | 5'-CTGCCAGTACTGCCGGCTAC-3'<br>5'-TCTGCACTGGAGTTCGTGGTATACT-3' |
| ABCA1 | 5'-GCACTGAGGAAGATGCTGAAA-3'<br>5'-AGTTCCTGGAAGGTCTTGTTCAC-3' |
| NPC1 | 5'-AGCCAGTAATGTCACCGAAAC-3'<br>5'-CCGAGGTTGAAGATAGTGTCG-3' |
| NPC2 | 5'-TATCCCTCTATAAAACTGGTGGTG-3'<br>5'-CCAGATGCACCGAACTCAAT-3' |
| RAB7A | 5'-CATCCTGGGAGATTCTGGAGTC-3'<br>5'-TGTGTCCCATATCTGCATTGTG-3' |
| ITGAX | 5'-GCTGAAGGCACACTGTGAAA-3'<br>5'-AGGGAGGCCGTGAAGTATCT-3' |
| CD44 | 5'-TGGCACCCGCTATGTCCAG-3'<br>5'-GTAGCAGGGATTCTGTCTG-3' |
| ILK | 5'-ATGTACTACATGAAGGCACCAATTTC-3'<br>5'-CCCCTTGCCATGTCCAAAG-3' |
| PADI3 | 5'-GGAGACCCTCGTGGACATTT-3'<br>5'-CTCCAAAGTCGCGTCAAAGC-3' |
| mGata2 | 5'-GCAGAGAAGCAAGGCTCGC-3'<br>5'-CAGTTGACACACTCCCGGC-3' |
| mGapdh | 5'-CTCCCACTCTTCCACCTTCG-3'<br>5'-GCCTCTCTTGCTCAGTGTCC-3' |
| | *PCR primers* |
| GATA2 | 5'-TATTTCCGGTGAATTCATGGAGGTGGCGGCCGAGCA-3'<br>5'-CGGGATCCGCGGCCGCTAGCCCATGGCGGTCACCATGCT-3' |
| | *shRNA sequences* |
| GATA2 | 5'-AAGGATCCAGCAAGGCTCGTTCCTGTTCATCAAG<br>AGTGAACAGGAACGAGCCTTGCTTTTTACCGGTAA-3' |
